# Weak seed banks influence the signature and detectability of selective sweeps

**DOI:** 10.1101/2022.04.26.489499

**Authors:** Kevin Korfmann, Diala Abu Awad, Aurélien Tellier

## Abstract

Seed banking (or dormancy) is a widespread bet-hedging strategy, generating a form of population overlap, which decreases the magnitude of genetic drift. The methodological complexity of integrating this trait implies it is ignored when developing tools to detect selective sweeps. But, as dormancy lengthens the ancestral recombination graph (ARG), increasing times to fixation, it can change the genomic signatures of selection. To detect genes under positive selection in seed banking species it is important to 1) determine whether the efficacy of selection is affected, and 2) predict the patterns of nucleotide diversity at and around positively selected alleles. We present the first tree sequence-based simulation program integrating a weak seed bank to examine the dynamics and genomic footprints of beneficial alleles in a finite population. We find that seed banking does not affect the probability of fixation and confirm expectations of increased times to fixation. We also confirm earlier findings that, for strong selection, the times to fixation are not scaled by the inbreeding effective population size in the presence of seed banks, but are shorter than would be expected. As seed banking increases the effective recombination rate, footprints of sweeps appear narrower around the selected sites and due to the scaling of the ARG are detectable for longer periods of time. The developed simulation tool can be used to predict the footprints of selection and draw statistical inference of past evolutionary events in plants, invertebrates, or fungi with seed banks.

## 1 Introduction

Seed banking is an ecological bet-hedging strategy, by which seeds or eggs lay in a dormant state of reduced metabolism until conditions are more favourable to hatch or germinate and complete the life-cycle. This life-history trait acts therefore as a buffer in uncertain environments (Cohen, 1966; Templeton and Levin, 1979) and has evolved several times independently in prokaryotes, fungi, plants, and invertebrates (Evans and Dennehy, 2005; Nara, 2009; Willis et al., 2014; Tellier, 2019; Lennon et al., 2021). Because several generations of seeds are simultaneously maintained, seed banks act as a temporal storage of genetic information (Evans and Dennehy, 2005), decreasing the effect of genetic drift and lengthening the time to fixation of neutral and selected alleles (Templeton and Levin, 1979; Hairston Jr and De Stasio Jr, 1988). Seed banks are therefore expected to play an important role in determining the adaptive potential of a species (Tellier, 2019). In bacteria (Shoemaker and Lennon, 2018; Lennon et al., 2021), invertebrates (Evans and Dennehy, 2005) or plants (Willis et al., 2014; Tellier, 2019), dormancy determines the neutral and selective diversity of populations by affecting the effective population size and buffering population size changes (Nunney and Ritland, 2002), affecting mutation rates (Levin, 1990; Whittle, 2006; Dann et al., 2017), genetic structure (Vitalis et al., 2004), rates of population extinction/recolonization (Brown and Kodric-Brown, 1977; Manna et al., 2017) and the efficacy of positive (Hairston Jr and De Stasio Jr, 1988; Koopmann et al., 2017; Heinrich et al., 2018; Shoemaker and Lennon, 2018) and balancing selection (Tellier and Brown, 2009; Verin and Tellier, 2018).

Seed banking, or dormancy, introduces a time delay between the changes in the active population and changes in the dormant population which considerably increases the time to reach the common ancestor of a sample of genes from the active population (Kaj et al., 2001; Blath et al., 2015, 2016, 2020). We note that two models of seed banks are proposed, namely the weak and strong dormancy models. These make different assumptions regarding the scale of the importance of dormancy relative to the evolutionary history of the species. On the one hand, the strong version is conceptualized after a modified two-island model with coalescence events occurring only in the active population as opposed to the dormant population (seed bank) with migration (dormancy and resuscitation) between the two (Blath et al., 2015, 2016, 2019; Shoemaker and Lennon, 2018). Strong seed bank applies more specifically to organisms, such as bacteria or viruses, which exhibit very quick multiplication cycles and can stay dormant for times on the order of the population size (thousands to millions of generations, Blath et al., 2015, 2020; Lennon et al., 2021). On the other hand, the weak seed bank model assumes that dormancy occurs only over a few generations (tens to hundreds), thus seemingly negligible when compared to the order of magnitude of the population size (Kaj et al., 2001; Tellier et al., 2011; Živković and Tellier, 2012; Sellinger et al., 2019), making it applicable to plant, fungi or invertebrate (*e*.*g. Daphnia sp*.) species or when the seed bank is experimentally imposed (as it is in practice difficult to generate the strong seed bank) (Shoemaker et al., 2022). We focus here on a pseudo-diploid version of the weak seed bank model in order to provide novel insights into the population genomic analysis of species which undergo sexual reproduction. The applicability of our results, as well as the differences and similarities between the strong and weak seed bank models, are highlighted in the Discussion.

The weak seed bank model can be formulated forward-in-time as an extension of the classic Wright-Fisher model for a population of size *N* haploid individuals. The constraint of choosing the parents of offspring at generation *t* only from the previous generation (*t −* 1) is lifted, and replaced with the option of choosing parents from previous generations (*t −* 2, *t −* 3, … up to a predetermined boundary *t − m*)(Nunney and Ritland, 2002). The equivalent backward-in-time model extends the classic Kingman coalescent and assumes an urn model in which lineages are thrown back-in-time into a sliding window of size *m* generations, representing the past populations of size *N* (Kaj et al., 2001). Coalescence events occur when two lineages randomly choose the same parent in the past. The germination probability of a seed of age *i* is *b*_*i*_, which is equivalent to the probability of one offspring choosing a parent *i* generations ago. The weak dormancy model is shown to converge to a standard Kingman coalescent with a scaled coalescence rate of 1*/β*, in which 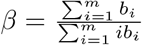 is the inverse of the mean time seeds spend in the seed bank, and *m* is the maximum time seeds can be dormant (Kaj et al., 2001). The intuition in a coalescence framework (Kaj et al., 2001) is that for two lineages to find a common ancestor, *i*.*e*. to coalesce, they need to choose the same parent in the active population, each the probability *β* to do so, as only active lineages can coalesce. Thus the probability that two lineages are simultaneously in the active population is a *β*^2^ scaling of the coalescence rate. The germination function was previously simplified by assuming that the distribution of the germination rate follows a truncated geometric function with rate *b*, so that *b* = *β* when *m* is large enough (Tellier et al., 2011; Živković and Tellier, 2012; Sellinger et al., 2019, see methods). A geometric germination function is also assumed in the forward-in-time diffusion model analysed in Koopmann et al., 2017; Heinrich et al., 2018 and Blath et al., 2020.

Seed banking influences neutral and selective processes via its influence on the rate of genetic drift. In a nutshell, a seed bank delays the time to fixation of a neutral allele and increases the inbreeding effective population size (from now on referred to only as the “effective population size”) by a factor 1*/b*^2^. The effective population size under a weak seed bank is defined as 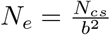 where *N*_*cs*_ is the census size of the active population (Nunney and Ritland, 2002; Tellier et al., 2011; Živković and Tellier, 2012). Mutation under an infinite site model can occur in seeds with probability *μ*_*d*_ and *μ*_*a*_ in the active population, so that we can define *θ* the population mutation rate under the weak seed bank model: 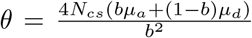 (Tellier et al., 2011). If mutations occur in the dormant population at the same rate as in the active population, we define *μ*_*d*_ = *μ*_*a*_ = *μ* yielding 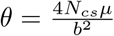, while if the dormant state does not mutate, *μ*_*d*_ = 0 and *μ*_*a*_ = *μ*, yielding 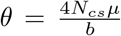. Empirical evidence (Levin, 1990; Whittle, 2006; Dann et al., 2017) and molecular biology experiments have shown that even under reduced metabolism DNA integrity has to be protected (Waterworth et al., 2016), and suggest that mutations occur in the dormant population (for simplicity at the same rate as in the active population, see model in Sellinger et al., 2019). Furthermore, recombination and the rate of crossing-over is also affected by seed banking. However, only the non-dormant lineages are affected by recombination in the backward-in-time model so that the population recombination rate is 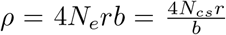. The recombination rate *r* needs to be multiplied by the probability of germination *b* as only active individuals can recombine (Živković and Tellier, 2018; Sellinger et al., 2019). The ratio of the population mutation rate and the recombination rate defines the amount of nucleotide diversity in the genome as well as the amount of linkage disequilibrium, a property which has been used to develop a sequential Markovian coalescent (SMC) approach to jointly estimate past demographic history and the germination rate (Sellinger et al., 2019, 2021).

While there is now a thorough understanding of how neutral diversity is affected by seed banking, the dynamics of alleles under selection have not been fully explored. Koopmann et al., 2017 developed a diffusion model of infinite (deterministic) seed bank model with positive selection and show that the time to fixation is not multiplied by 1*/b*^2^ (as for neutral alleles) but by a higher factor (between 1*/b*^2^ and 1*/b*). The interpretation is as follows: while the time to fixation of an advantageous allele is lengthened compared to a model without dormancy, the efficacy of selection should be altered compared to a neutral allele (the effect of genetic drift). Namely, the Site Frequency Spectrum (SFS) of independently selected alleles shows an increased deviation from neutrality with a decreasing value of *b*. By relaxing the deterministic seed bank assumption, Heinrich et al., 2018 find that: 1) a finite small seed bank decreases the efficacy of selection, and 2) selection on fecundity (production of offspring/seeds) yields a different selection efficiency compared to selection on viability (seed viability), as can be seen from their estimated Site-Frequency Spectrum (SFS) of independent alleles under selection. Furthermore, based on the effect of seed banks on *θ* and *ρ* and on selection, verbal predictions on the genomic signatures of selection have been put forth (Živković and Tellier, 2018).

These theoretical and conceptual approaches, while paving the way for studying selection under seed banks, did not consider the following argument. If the time to fixation of an advantageous allele increases due to the seed bank, it can be expected that 1) drift has more time to drive this allele to extinction, and 2) the signatures of selective sweeps can be erased by new mutations appearing in the vicinity of the selected alleles. These effects would counter-act Koopmann et al.’s (2017) predictions that selection is more efficient under a stronger seed bank compared to genetic drift, as well as Živković and Tellier’s (2018), that selective sweeps are more easily observable under stronger seed bank. In order to resolve this paradox, we develop and make available the first simulation method for the weak seed bank model, which allows users to generate full genome data under neutrality and selection. We first present the simulation model, which we use to follow the frequencies of an adaptive allele in a population with seed banking. We aim to provide insights into the characteristics of selective sweeps, including the time and probability of fixation, as well as recommendations for their detection in species exhibiting seed banks.

## 2 Methods

Forward-in-time individual-based simulations are implemented in C++. Genealogies are stored and manipulated with the tree sequence toolkit (tskit, Kelleher et al., 2018), which allows for a general approach to handling arbitrary evolutionary models and an efficient workflow through well-documented functions.

### 2.1 Model

The model represents a single, panmictic population of *N* hermaphroditic pseudo-diploid adults, which will henceforth be referred to as diploids for brevity. Population size is fixed and generations are discrete, so that in the absence of dormancy and selection, the population follows a classic Wright-Fisher model. In this case, at the beginning of each generation, a new individual is produced by sampling two parents from the previous generation. Once sampled, each parent contributes a (recombined) gamete to generate the new individual. Each parent is sampled with probability 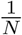 (multinomial sampling), leading to two vectors **X**_*parent*1_ and **X**_*parent*2_, containing the indices of the respective parents:

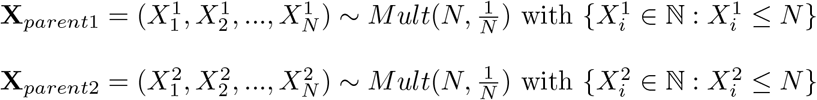

Dormancy adds a layer of complexity, by introducing seeds that can germinate after being dormant for many generations. This relaxes the implicit Wright-Fisher assumption, as parents are no longer only sampled from the previous generation, but also from dormant individuals produced up to *m* generations in the past. The probability of being sampled from generation *k* depends on the probability of germination, which is a function of the age of the dormant individual. As for the classical Wright-Fisher model, there are 2*N* possible parents. The parents are sampled using a probability vector **Y**^*norm*^ written as:

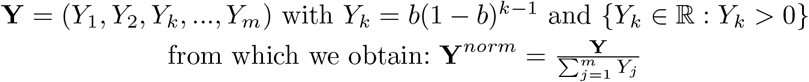

From the expression above, the probability of being sampled follows a truncated geometric distribution parameterized with germination rate *b* and then normalized. The generation *G* of each parent is randomly sampled using a multinomial sampling with the probability vector **Y**^*norm*^.

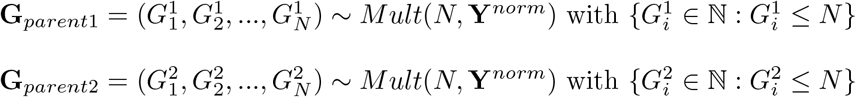

Once the age of each of the 2*N* parents has been determined, random individuals from the corresponding age groups are sampled (the same individual can be sampled more than once) and one recombined gamete from each of these 2*N* individuals is generated. These gametes are then randomly combined to form *N* new diploid individuals which constitute the current active population. Thus, the forward simulation process models two haploid dormant individuals (with different ages) which become active at the current generation and join to form a diploid individual (Figure 1). This pseudo-diploid model formulation is implicitly equivalent to haploid gametes being resuscitated from the dormant state and fusing to create a diploid individual capable of reproduction. The probability of coalescence (*p*_*coal*_) is therefore expected to follow haploid expectations 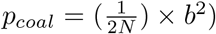.

**Fig. 1.**
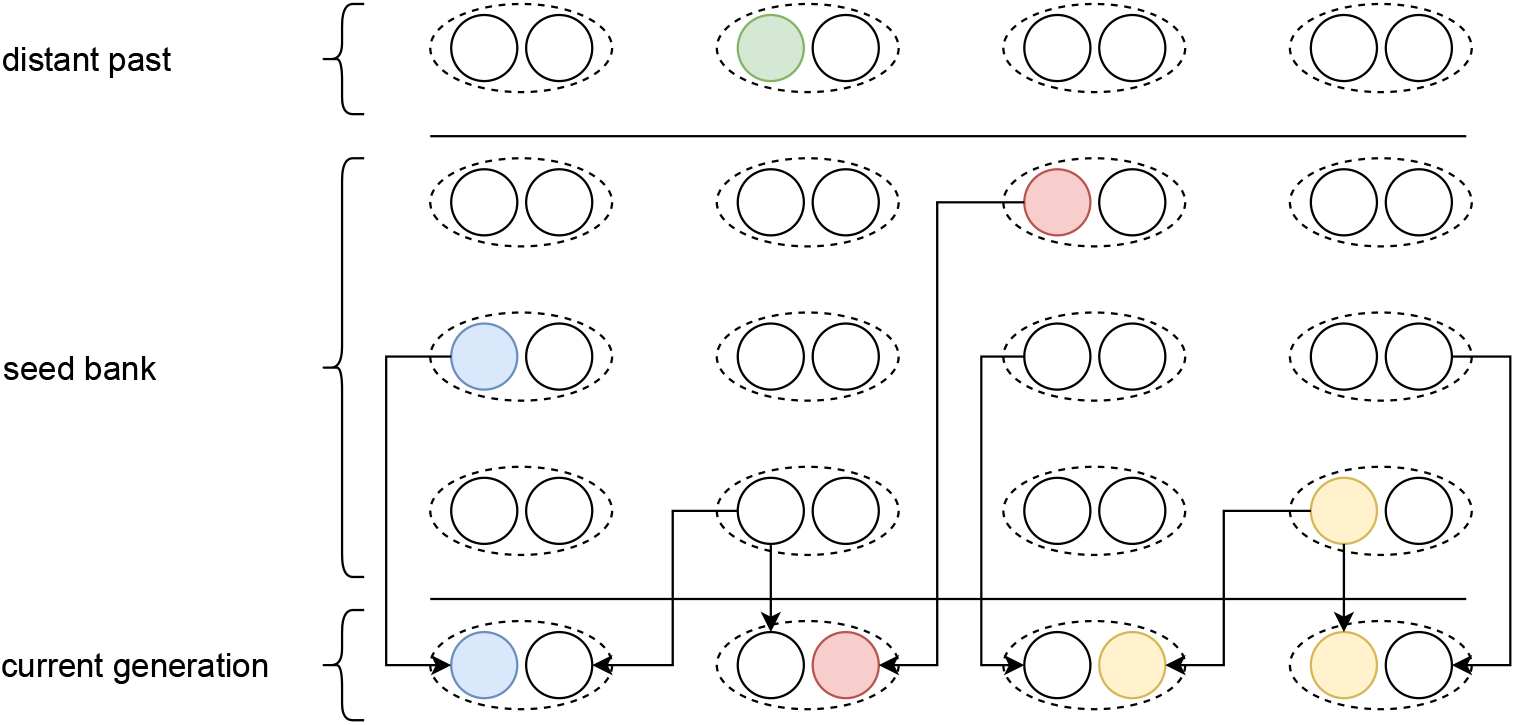
Schematic representation of our pseudo-diploid weak dormancy seed bank model by a forward-in-time two step process in the spirit of Kaj et al., 2001 for haploid dormant seeds. The arrows originating from the parent or seed generation represent the geometric sampling process of the current generation, and the sampling of the individual within the given generation of the past based on the respective fitness value.

The number of recombination events is sampled from a Poisson distribution with parameter *r* (for example 1 *×* 10^*−*8^ per bp per generation). At the end of this process, new mutations can be introduced (only necessary for sweep detection tools). Generally neutral mutations are not simulated and statistics are computed using branch lengths. We assume here that mutations are also introduced at every generation in dormant individuals at the same rate (following Sellinger et al., 2019), even if they are not explicitly simulated. Recombination breakpoints are uniformly distributed across the genome with each coalescent tree being delineated by two recombination breakpoints.

To model selection signatures within a neutral genomic background, we consider non-neutral bi-allelic loci, placed at predefined and fixed genomic positions, with beneficial mutations arising after the burn-in period. A locus under selection has a dominance *h* and selection coefficient *s*, respectively. The expressions for the fitness of heterozygote and homozygote individuals with the beneficial mutation are thus 1 + *hs* and 1 + *s*, respectively. Fitness affects the probability that an individual’s gametes can leave the dormant state and contribute to reproduction. The choice of the germinating generation when sampling the parents is unaffected by their fitness values, but the sampling of individuals within a given generation is determined by the fitness. In other words, selection acts on fecundity, as the fitness of an allele determines the number of offspring produced and not survival (viability selection). A selection coefficient of 0 would lead to multinomial Wright-Fisher sampling, which can be used to track neutral mutations over time. This two-step process of first choosing the generation followed by the individual is presented in Figure 1.

From a technical perspective, individuals can be tracked in the tskit-provided table data structures, if the *tree_sequence_recording* feature is enabled. This feature is not required when computing statistics on allele frequency dynamics only (*i*.*e*. to compute fixation times or probabilities). The tables used in this simulation are as follows: 1) a node table representing a set of genomes, 2) an edge-table defining parent-offspring relationships between node pairs over a genomic interval, 3) a site table to store the ancestral states of positions in the genome, and 4) a mutation table defining state changes at particular sites. The last two tables are only used to introduce the mutation under selection. If neutral mutations are required for down-stream analysis, they are simulated after this step. The simulation code works with the aforementioned tables through tskit functions, e.g. the addition of information to a table after sampling a particular individual or through the removal of parents who do not have offspring in the current generation in a recurrent simplification process. This clean-up process is a requirement to reduce RAM-usage during the simulation, because keeping track of every individual ever simulated to build the genealogy quickly becomes infeasible. However, a noticeable difference to the classic use of the tskit function is that in our case individuals which have not produced offspring in the past, but are still within the dormancy upper-bound defined range of *m* generations, need to be protected from the simplification process, which is achieved by marking them as *sample nodes* during the simulation. Indeed, forward-in-time, a parent can give offspring many generations later (maximum *m*) through germinating seeds. As previously stated, the simulation process can be run independently of tskit, but the latter is required when planning to analyze the genealogy.

### 2.2 Simulations

Except when indicated otherwise, the population size is generally set to *N* = 500 individuals or 2*N* = 1, 000 haploid genomes. We specifically change population size when testing whether sweep signatures can be explained by simple size scaling. In this case we use *N* = 2000 individuals with a germination rate of *b* = 1, corresponding to *N* = 245 for *b* = 0.35 (Figure

The genome sequence length is set to 100,000 bp, 1MB or 10 MB. Simulations start with a burn in or calibration phase of 50,000 generations for *b* = 1, and 200,000 generations for *b* = 0.5 (see Figure

Neutral diversity is calculated based on the branch length, meaning that explicitly simulating mutations is not required. To check whether the strength of a sweep behaves in accordance to expectations *i*.*e*. lower recombination rates result in wider sweeps, recombination rates ranging from 5 *×* 10^*−*8^ to *r* = 10^*−*7^ are tested for all parameter sets. Simulations are run for the germination rate *b* ranging from 0.25 up to 1 (with *b* = 1 meaning no dormancy). The upper-bound number of generations *m* which is the maximum time that seeds can remain dormant (*i*.*e*. seeds older than *m* are removed from the population) is set at 30 generations. Beneficial mutations have a selective coefficient 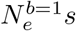 ranging from 0.1 to 100 and dominance *h* takes values 0.1, 0.5 and 1.1, representing recessive, co-dominant and overdominant beneficial mutations.

### 2.3 Statistics and sweep detection

We first calculate several statistics relative to the forward-in-time change of the frequency of an advantageous allele in the population, such as the mean time to fixation and the probability of fixation, using 1,000 simulations per parameter configuration. Each simulation run consists of the recurrent introduction over time of an allele (mutant at frequency 1*/*2*N*) which is either lost or fixed. When an allele is lost and the simulation is conditioned on fixation a new simulation starts from a neutral genetic diversity background (see below for more details). An allele is considered to be fixed if its number of copies is 2N for *m* consecutive generations. For each simulation run we store 1) the time it takes for the last introduced allele to reach fixation (time between allele introduction until fixation), and 2) the number of alleles which were introduced until one has reached fixation (yielding the probability of fixation of an allele per simulation run). The resulting times to fixation and fixation probabilities are calculated as the averages over the 1,000 simulation runs.

We also compute statistics on the underlying coalescent tree and ancestral recombination graph (ARG) such as time to the most recent common ancestor, linkage disequilibrium (*r*^2^, Hill and Robertson, 1968), as well as Tajima’s *π* and D (Tajima, 1983; Nei and Li, 1979; Tajima, 1989) over windows of size 5,000 (giving 200 windows for a sequence length of 1 MB). This allows us to analyse the effects of seed-dormancy on the amount of linkage disequilibrium and nucleotide diversity along the genome, as well as the footprint of a selective sweep on these quantities. *Tskit* functions are used for diversity and linkage disequilibrium calculations. Nucleotide diversity (*π*) is calculated based on the branch length. Sweeps are detected using Omega and SweeD statistic, the first one quantifies the degree to which LD is elevated on both sides of the selective sweeps, as implemented and applied with OmegaPlus (Alachiotis et al., 2012), while SweeD (Pavlidis et al., 2013) uses changes in SFS across windows to detect sweeps. A difficult issue in detecting selective sweeps is choosing the correct window size to perform the computations. It is documented that the optimal window size depends on the recombination rate and thus the observed amount of linkage disequilibrium (Alachiotis et al., 2012; Alachiotis and Pavlidis, 2016). We use two different setups with different window sizes: – minwin 2000 –maxwin 50000 and –minwin 1000 –maxwin 25000. The window sizes refer to the minimum and maximum region used to calculate LD values between mutations. Importantly the –minwin parameter determines the sensitivity, meaning the degree to which false positives or false negatives (high –minwin values) are detected, while the –maxwin parameter determines run-time and memory requirements. A detailed graphical description can be found in the online OmegaPlus manual. In theory the larger window size is more appropriate for the model without dormancy (*b* = 1), and the narrower window size for the model with dormancy (*b <* 1). For both cases, we set –grid 1000 –length 10 MB. SweeD is only tested using a –grid 1000 parameter. The statistic is computed for a sample size of 100 over 400 simulations for each sweep signature at mulitple generations after fixation (sweep recovery scenerios).

### 2.4 Code description and availability

Source code of the simulator and demonstration of the analysis can be found at https://gitlab.lrz.de/kevin.korfmann/sleepy and https://gitlab.lrz.de/kevin.korfmann/sleepy-analysis. A convenient feature of the simulator is the option to choose between switching the tree sequence recording on or off depending on the question, *i*.*e*. if analysing fixation time and probability of fixation it is unnecessary to record the tree sequence (or use a calibration phase). To analyse the sweep signatures, the simulation process has been divided into two phases to alleviate the large run-times of forward simulations. During the first phase, a tree sequence will be generated under neutrality and stored to disk. And in the second phase the neutral tree sequence is loaded and a parameter of interest is tested until fixation or loss. Additionally, if the simulation is conditioned on fixation, then the simulation can start again from the beginning of the second phase that will have been run for tree sequence calibration, saving the time.

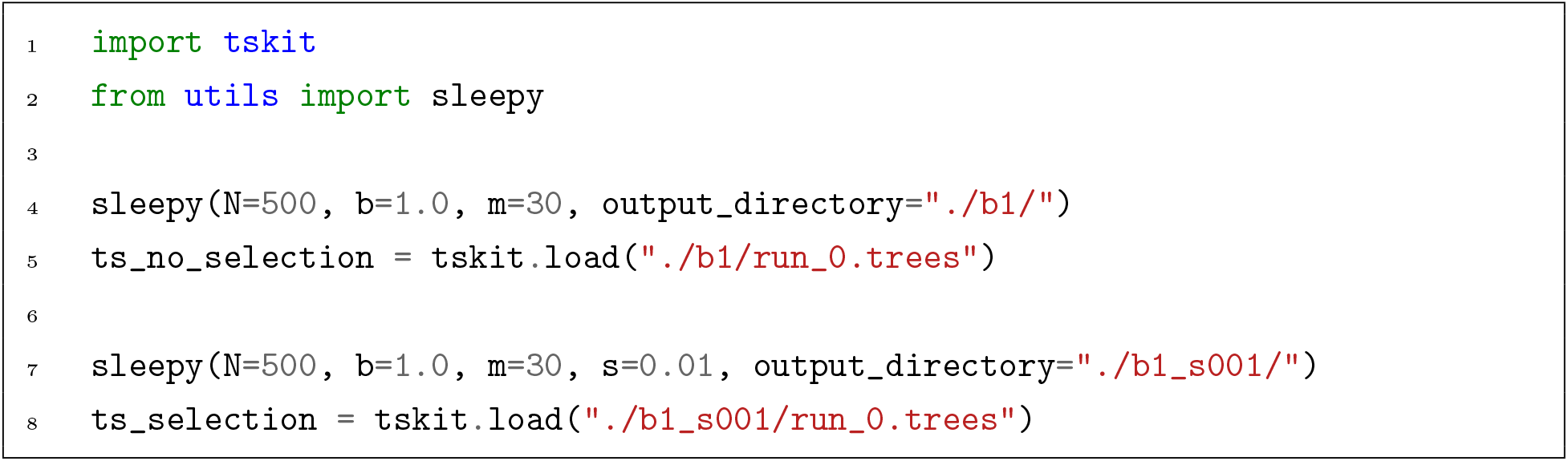

Listing 1: Simplified, demonstrative Python code example for a simulation with and without selection. Tree sequence results are stored in a specified output directory and are loaded via *tskit* function for further processing or analysis of e.g. linkage disequilibrium or nucleotide diversity along the genome. A more detailed version with more parameters can be found in the example notebook at https://gitlab.lrz.de/kevin.korfmann/sleepy-analysis.

Simulations rely on regular simplification intervals for efficiency of the genealogy recording, yet the weak dormancy model requires keeping up to *m* generations in memory even for past individuals (seeds) which do not have offspring in the current generation. To make sure that this assumption is realized in the code, up to *m* generations are technically defined as leaf nodes, thus hiding them from the regular memory clean-up process. Furthermore, the presence or absence of an allele with an associated selection coefficient needs to be retrievable, even under the influence of recombination, for all individuals for up to *m* generations in order to determine the fitness of the potential parents. Therefore, recombination and selective alleles are tracked additionally outside of the *tskit* table data structure, allowing the running of the the simulation without the tree sequence. Both of these model requirements, namely maintaining individuals which do not have offspring in the current generation (but potentially could have due to stochastic resuscitation of a seed) as well as the knowledge about the precise state of that given individual in the past, are reasons to choose our own implementation over SLiM (Haller and Messer, 2019).

## 3 Results

### 3.1 Neutral coalescence

We first verify that our simulator accurately produces the expected coalescent tree in a population with a seed bank with germination parameter *b* and population size *N*. To do so, we first compute the time to the most recent common ancestor (TMRCA) of a coalescent tree for a sample size *n* = 500.

We find that the coalescent trees are scaled by a factor 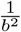 independently of the chosen recombination rate (Figure 2a). The variance of the TMRCA decreases with increasing recombination due to lower linkage disequilibrium among adjacent loci, as expected under the classic Kingman coalescent with recombination (Hudson, 1983). Moreover, we also find that decreasing the value of *b* (*i*.*e*. maintaining the dormant population for longer) decreases linkage disequilibrium (Figure 2b). This is a direct consequence of the scaling of the recombination rate by 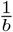, because any active individual can undergo recombination (and can be picked as a parent with a probability *b* backwards in time). Therefore, we observe here two simultaneous effects of seed banks on the ARG: 1) the length of the coalescent tree and the time between coalescence events is increased by a factor 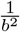 meaning an increase in nucleotide diversity (under a given mutation parameter *μ*), and 2) a given lineage has a probability *br* to undergo an event of recombination backward in time. In other words, even if the recombination rate *r* is slowed down by a factor *b* (because only active individuals recombine), since the coalescent tree is lengthened by a factor 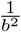 there are on average 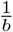 more recombination events per chromosome.

**Fig. 2.**
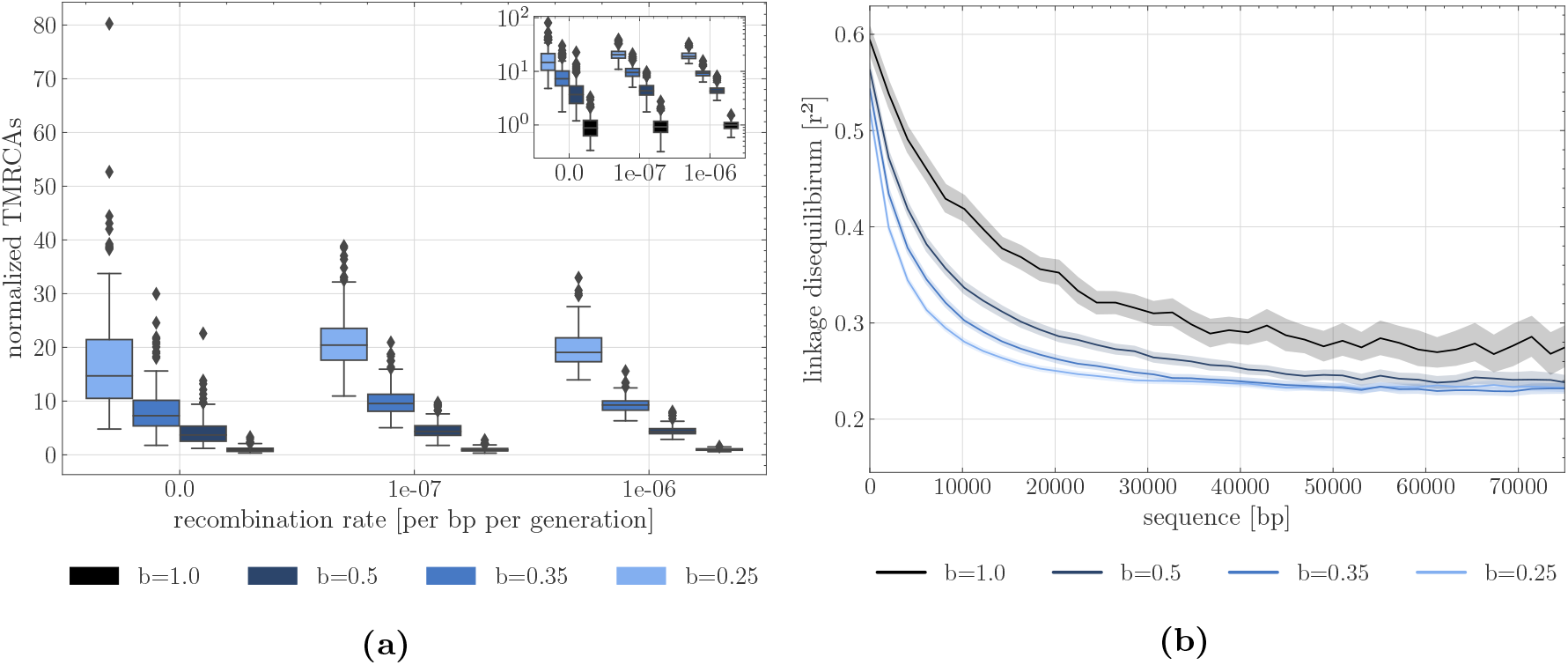
(a) Time to the most recent common ancestor (TMRCA) as a function of the germination rate *b* and scaled by results under *b* = 1. For each germination rate, three recombination rates per site are presented (*r* = 0, *r* = 10^*−*7^ and *r* = 10^*−*6^. Boxes describe the 25th (Q1) to 75th percentile (Q3), with the lower whisker representing Q1-1.5 *×* (Q3-Q1) outlier threshold and the upper whisker is calculated analogously. The mean is plotted between Q3 and Q1. Each boxplot represents the distribution of 200 TMRCA values over 200 sequences of 0.1 Mb. Per sequence the oldest TMRCA is retained. (b) Monotonous decrease of linkage disequilibrium as a function of distance between pairs of SNPs, setting *r* = 10^*−*7^ per generation per bp, sequence length to 10^5^ bp. While population size is 500, linkage decay was calculated by subsetting 200 individuals, purely to constrain the computational burden. In total 200 replicates were used for TMRCA and LD calculations. Shaded areas represent the 95 % confidence interval.

This property of the ARG was used in Sellinger et al., 2019 to estimate the germination parameter using the sequential Markovian coalescent approximation along the genome.

### 3.2 Allele fixation under positive selection

We examine the trajectory of allele frequency of neutral and beneficial mutations, by computing the probabilities and times to fixation over 1000 simulations. As expected for the case without dormancy (*b* = 1), the probability of fixation of a beneficial allele increases with the strength of selection (Figure 3a).

**Fig. 3.**
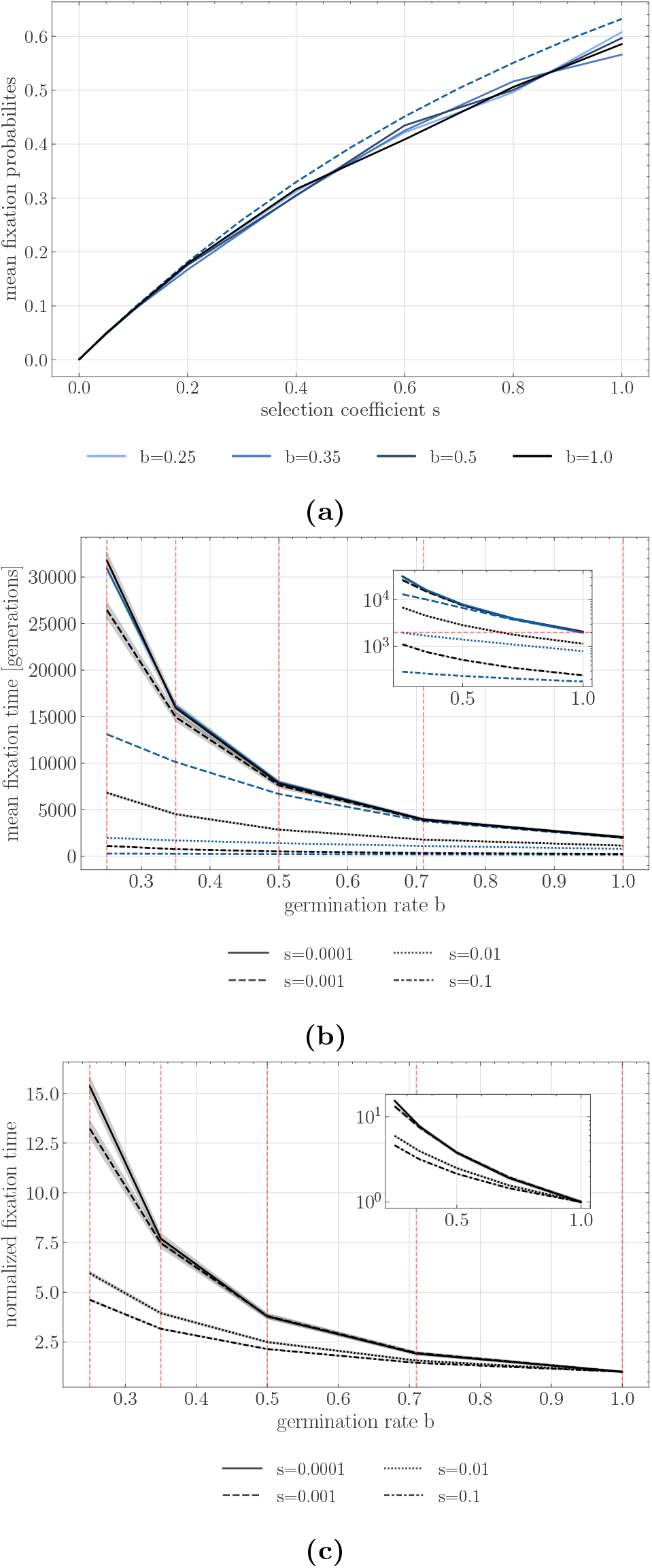
(a) Simulated estimates of the probability of fixation of an advantageous allele with different coefficients of selection *s* under absence of seed bank *b* = 1 (black solid line) and various seed bank strength *b* = 0.5, 0.35, 0.25 (blue lines) along with the theoretical expectations for a neutral allele (dashed). (b) Time to fixation for different selection coefficients. Y-axis is the time in generations, and X-axis is the germination rate *b*. (c) Normalized time to fixation with respect to the number of generations for *b* = 1 for each selection coefficient version of b). In b) and c) black lines represent time to fixation under seed bank. The blue lines indicate the time to fixation in a population with-out dormancy but with an effective population size scaled by 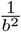 and the re-spective scaled effective selection coefficient 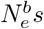. For example, for *s* = 0.001, we quantify the fixation time of alleles under 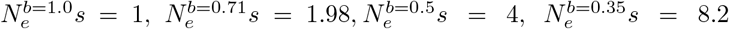, and 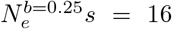, and (indicated by the red vertical dashed lines). Population size is 500 diploids, *h* = 0.5, 1,000 replicates are used for each parameter combination, and shaded areas represent the 95% confidence interval.

We note, that the mean fixation probability is unaffected by the seed bank, as when *N*_*e*_ is large enough and the coefficient of selection *s* is not too strong, the probability of fixation of a beneficial mutation depends only on *hs* (Barrett et al., 2006).

As expected from the neutral case, the time to fixation with dormancy becomes longer with smaller values of *b* (Figure 3b). When selection is weak the time to fixation is close to the expectation for neutral mutations (Figure 3b, *b* = 1: 4*N* = 2000 generations and 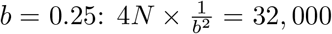 generations). However, increasing *s* changes the scaling of the time to fixation. Dormancy significantly increases the times to fixation, beyond that expected by *N*_*e*_. This can be seen by comparing the expectations for the times to fixation for the rescaled effective population size without dormancy (blue lines in 3b) to those obtained from our simulations (black lines). In order to understand this observation, we examine the time an allele under selection remains at given frequencies in the active population. The trajectory of an allele undergoing selection can be separated into three phases: two that are qualified as “stochastic”, when the allele is at a very low or very high frequency, and one “deterministic”, during which the frequency of the allele increases exponentially (see Kim and Stephan, 2002). As shown in Figures S2-4, we find that the proportion of time spent at very low and very high frequencies increases with increasing selection and decreasing *b* (it is unaffected by *b* when selection is weak *i*.*e. s* = 0.0001). This observation, along with generally shorter relative times spent in the deterministic phase (Figure S4) with increasing *b*, imply that the seed bank contributes to increasing the duration of the stochastic phases, slowing down the selection process.

### 3.3 Footprints of selective sweep

Now that we have a clearer indication of the dynamics of allele fixation, we use our new simulation tool to investigate the genomic diversity and signatures of selective sweeps at and near the locus under positive selection by simulating long portions of the genome (Figure 4). In accordance with the results from Figures 2a and 2b and the effects of the seed bank in maintaining genetic diversity, smaller germination rates lead to higher neutral genetic diversity due to the lengthening of the coalescent trees (e.g. Figure 4a measured as Tajima’s *π*). Moreover, comparing the width of the selective sweeps valley of polymorphism in presence and absence of dormancy, we conclude that stronger dormancy generates narrower selective sweeps around sites under positive selection which have reached fixation (Figures 4b, 4d and

**Fig. 4.**
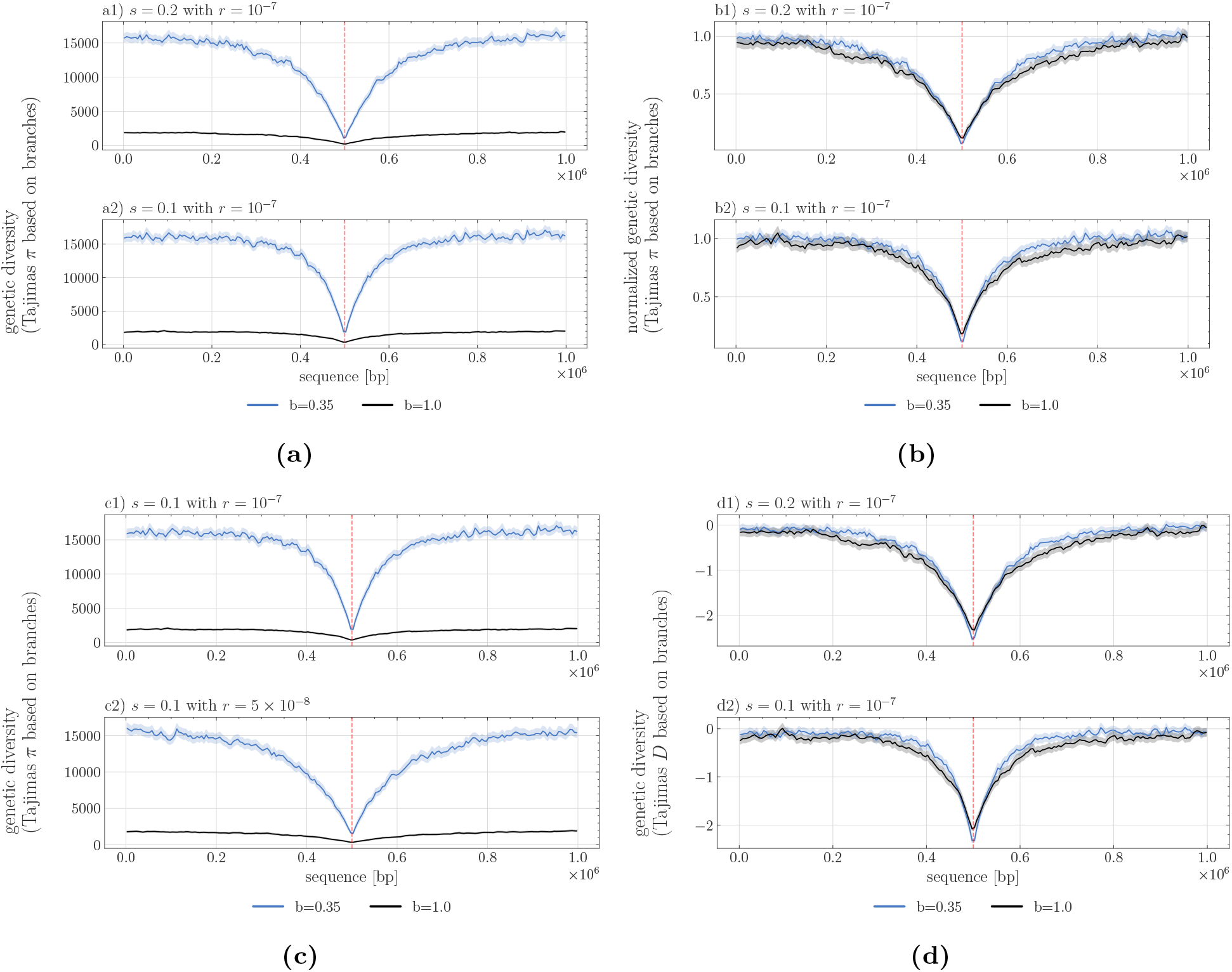
Signature of selective sweeps as measured by nucleotide diversity (Tajimas *π* in a, b, c) and Tajimas D (in d) over 1Mb sequence length (X-axis), the selected site being located in the middle of the segment. The statistics are computed per windows of size 5,000 bp and averaged over 200 repetitions, the shaded area representing the 95% confidence interval. The black line indicates the value without a seed bank (*b* = 1) and the blue line with dormancy (*b* = 0.35). a) *π* assuming two selection coefficients 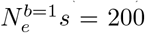 (a1) and 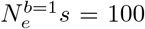 (a2) with *h* = 0.5. (b) Normalized *π* as divided by the average neutral branch diversity, namely approx. 2000 for *b* = 1 and approx. 16000 for *b* = 0.35 (see (a) or (c) between sequence range of 0 to 0.2 *×* 10^6^ or from 0.8 *×* 10^6^ to 1 *×* 10^6^). (c) *π* assuming two recombination rates *r* = 10^*−*7^ per bp per generation (c1) and *r* = 5 *×* 10^*−*8^ per bp per generation (c2).

### 3.4 Detectability of selective sweeps

Based on the previous results, we hypothesize that, compared to the absence of seed banking, the detectability of selective sweeps in a species with seed bank is affected 1) in the genome space, that is the ability to detect the site under selection, and 2) in time, that is the ability to detect a sweep after the fixation of the beneficial allele. First, as the footprints of selective sweeps are sharper and narrower in the genome under a stronger seed bank, we expect that the detection of these sweeps likely requires adapting the different parameters of sweep detection tools, namely the window size to compute sweep statistics. Second, in a population without dormancy, the time for which the detection of a selective sweep signature is possible is approximately 0.1N generations (Kim and Stephan, 2002). We hypothesize that as the mutation rate and genetic drift are scaled by 1*/b*^2^, the time it takes a sweep to recover after it has reached the state of fixation is slowed down. The time window for which a sweep could still be detected would then be potentially longer than 0.1*N* generations.

In Figure 5 we show the results obtained using OmegaPlus and SweeD, both tools for detecting selective sweeps (Alachiotis et al., 2012; Pavlidis et al., 2013). As noted above, individual simulations show significant variation in nucleotide diversity and LD, which is not captured by the mean diversity over several runs plotted in the figures above. As the detection of sweeps is performed against the genomic background of each individual simulation, these variations in nucleotide diversity and LD generate confounding effects and define the rates of false positives expected from the detection test. Following the classic procedure to detect sweeps, we use neutral simulations to define different thresholds for detection, for which we obtain a false positive rate of less than 0.05. We find that when using the same large detection window “–minwin 2000 –maxwin 50000” for *b* = 1 and *b* = 0.35 (Figures 5 a21 and 5 b21), sweep detection almost completely fails for *b* = 1, unless the fixation has just occurred, meaning that no generation has passed since the fixation event. For *b* = 0.35 sweeps are detectable up to *>*2000 generations after fixation. Decreasing the window size is generally associated with a loss of sensitivity, increasing the rate of true and false positives. This is true for *b* = 1 (see neutral threshold line in Figure 5 b21 and b22), indicating a decrease from roughly 60 % detected sweeps to 40 % (after 400 repetitions). However, the detectability of older sweeps (*>*2,000 generations) is increased for *b* = 0.35 (Figure 5 b22). Results using SweeD support this increased detectability, also when using the SFS statistics, showing the possibility of locating sweeps approximately up to 2,000 generations after fixation (Figure 5 a3 and b3)

**Fig. 5.**
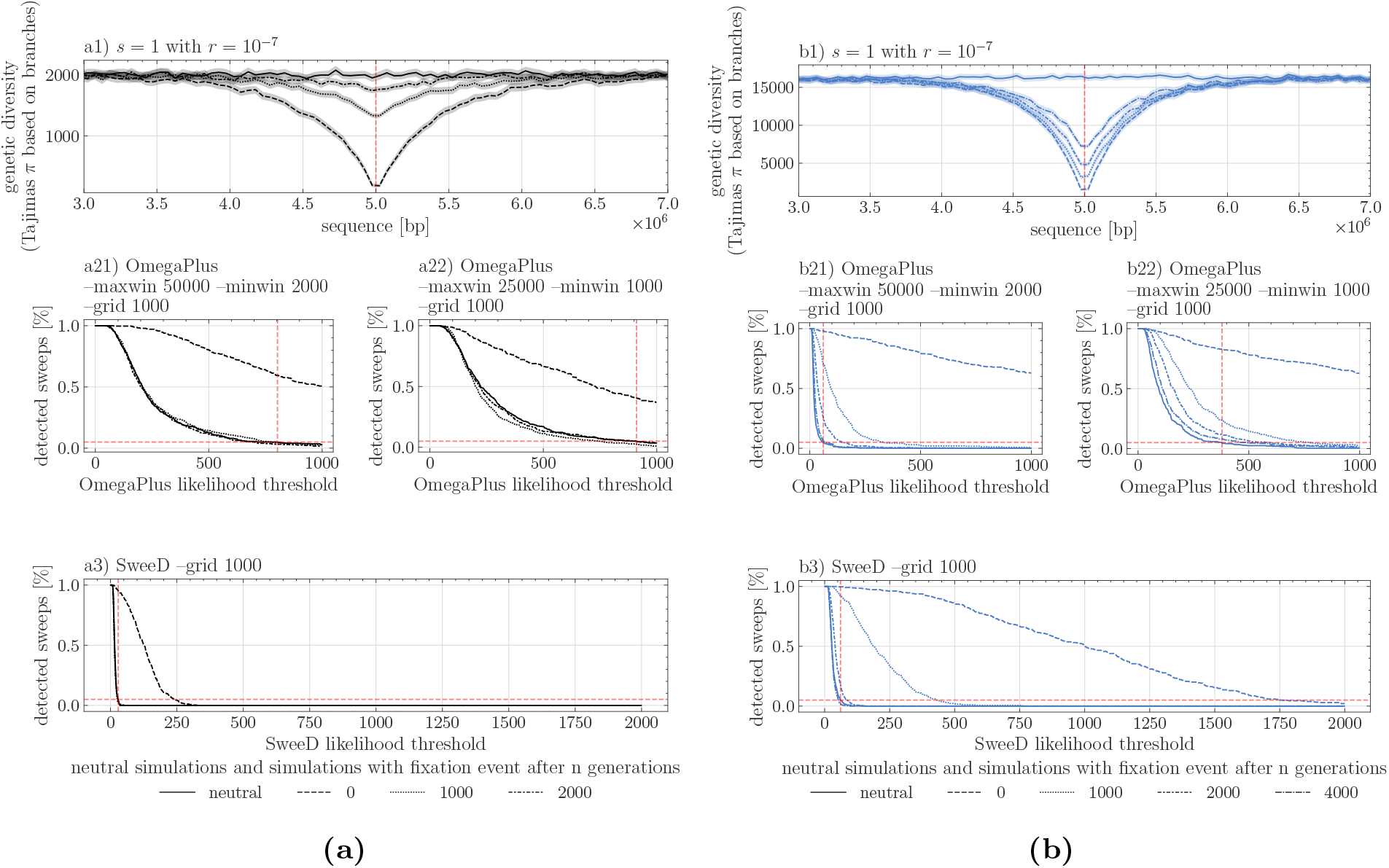
Selective sweep detection depending on the threshold of OmegaPlus or SweeD statistics on a 10MB sequence with a strong selective mutation of 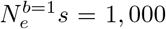 located in the middle of the sequence. Two germination rates apply: a1) *b* = 1 and b1) *b* = 0.35, with the signature of sweep being shown at various time points after the fixation event (0, 1000, 2000 and 4000 generations). Results for two window sizes “–minwin 2000 –maxwin 50000” (a21,b21) and “–minwin 1000 –maxwin 25000” (a22,b22) for analysis with OmegaPlus and SweeD (a3 and b3) using a grid size of 1,000. The percentage of detected sweeps is indicated for a given user-defined threshold value on the X-axis. Vertical dashed lines indicate the 5% sweep detection based on neutral simulations, setting up the false positive rate. Recombination rate is *r* = 1 *×* 10^*−*7^ per bp per generation for all sweep simulations, and 400 replicates for each parameter.

We note that there is a much sharper decrease in the rate of detection of false positive sweeps (neutral simulation line in Figure 5) under seed bank compared to the absence of a seed bank, likely being a direct consequence of the increased linkage decay around the site. Lastly, the possibility to locate sweeps multiple generations after the fixation event emphasizes the slower recovery of nucleotide diversity post-fixation in combination with the already established narrowness of the signature in the presence of a seed bank for a given population size *N* (*b* = 0.35, Figure

## 4 Discussion

We investigate the neutral and selective genome-wide characteristics of a weak seed bank model by means of a newly developed simulator. We first characterize the emergent behavior of an adaptive allele under a weak seed bank model, and simulate the times to and probabilities of fixation, considering different strengths of selection and recombination. In populations without seed banks, a neutral mutation is expected to fix after a time of 2*N*_*e*_ generations and *≈* 2*N*_*e*_*s* if the allele is under weak selection (Kimura, 1962). Though both processes are re-scaled by the weak dormancy model (Koopmann et al., 2017), the time to fixation of a neutral mutation can be obtained by rescaling *N*_*e*_ appropriately (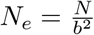 in the case of a seed bank, with *b* the germination rate). This remains true under weak selection, however under strong selection the time to fixation is significantly decreased and cannot be explained by the change in *N*_*e*_ alone. In accordance with existing theory, the probability of fixation is unaffected by the seed bank (since it depends only on *sh*, see for example Barrett et al., 2006), implying that the main effect of seed banks is on the dynamics of allelic frequencies, but not on the outcome of selection at a single locus. Combining this observation and the effect of seed banks on increasing the effective recombination rate, we suggest that the signatures of sweeps may be slightly easier to detect in the presence of seed banking as shown by the sharpness and depth of the nucleotide diversity pattern (the so-called valley of polymorphism due to genetic hitch-hiking, Maynard Smith and Haigh, 1974; Kim and Stephan, 2002) against the genomic background.

### 4.1 Dynamics of alleles under positive selection

Our results regarding the time to fixation of advantageous alleles are in line with previous works in showing that a weak seed bank delays the time to fixation (Hairston Jr and De Stasio Jr, 1988; Koopmann et al., 2017; Heinrich et al., 2018; Shoemaker and Lennon, 2018). However, a novelty here is that we refine these results in showing that the time to fixation of a weakly (*s <* 0.01) and a strongly (*s ≥* 0.01) positively selected allele differ under seed bank: the selection on weak alleles is delayed by a factor 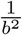 while for strong selection, the time to fixation is delayed by more than would be expected for a population without a seed bank but the same effective population size(see Figure 3b,3c, and Koopmann et al. 2017 for an analytical approach with an infinite deterministic seed bank). We show that this delay can be explained by an increase in the time spent in the stochastic phases of allele fixation (at below 10% and above 90% in the active population). In other words, dormancy delays the action of selection under the weak seed bank model (due to the dormant population acting as a buffer slowing down allele frequency change). In the initial phase of selection when the advantageous allele is at a very low frequency in the (active) population, and before reaching the exponential phase, the allele frequency increases almost deterministicly (Kim and Stephan, 2002). This delay in the initial selection phase is visible in Figure 4a in Shoemaker and Lennon, 2018. Our results are valid for the weak seed bank model (as studied in Figure 4a in Shoemaker and Lennon, 2018, and Koopmann et al., 2017) and we find that there exists a unique phase of selection encompassing the time until all individuals (in the active and dormant population) have fixed the advantageous allele. Strong seed bank models behave differently with respect to time to fixation of alleles under selection (Shoemaker and Lennon, 2018), showing two distinct phases: a first rapid phase of selection in the active population, followed by a second long delay until there is fixation in the dormant population. We are not aware of any results regarding the effect of strong seed banking on the probability of allele fixation. Our results thus mitigate the previous claim that (weak) seed banks may amplify selection, making it relatively more efficient with regards to the effects of genetic drift, while it does not alter the probability of fixation of an advantageous allele.

Longer times to fixation should promote genetic diversity, but as the probability of fixation at a single locus is unchanged by the seed bank, dormancy does not necessarily enhance the adaptive potential (by positive selection) of a population.

### 4.2 Signals of selective sweeps

The precise signature of a positive selective sweep is dependent on a variety of factors, *i*.*e*. age of the observation after fixation, degree of linkage due to recombination, and its detectability depends on the specified window size to compute polymorphism statistics. However, in the case of sweeps under seed bank, two effects are at play and change the classic expectations based on the hitch-hiking model without generation overlap. First, as the effective population size under seed bank increases with smaller values of *b*, an excess of new mutations is expected to occur after fixation around the site under selection compared to the absence of seed bank. As these new mutations are singleton SNPs, we suggest that the signature of selective sweeps observed in the site-frequency spectrum (U-shaped SFS) should be detectable under seed bank (Maynard Smith and Haigh, 1974; Kim and Stephan, 2002). Additionally, this effect was also detectable by the other sweep detection methods based on the SFS (SweeD, Pavlidis et al., 2013), finding sweeps older than 2000 generations (for N=500).

Second, the signature of sweeps also depends on the distribution of linkage disequilibrium (LD) around the site under selection (Alachiotis et al., 2012; Bisschop et al., 2021), which is affected by the seed bank (Figure 4). Theoretically, it has been shown that patterns of LD both on either side and across the selected site generally provide good predictive power to detect the allele under selection. We use this property when using OmegaPlus, which relies on LD patterns across sites. Further past demography should be accounted to correct for false positives, due for example to bottlenecks (see review in Stephan, 2019). We speculate that a high effective recombination rate around the site under selection, as a consequence of the seed bank, maybe an advantage when detecting sweeps. This allows the avoidance of confounding effects due to the SFS shape, which is sensitive to demographic history. We also highlight that the narrower shape of the selective sweep under stronger seed bank, and the smaller number of loci contained in the window, reduce the number of false positives.

As mentioned above, a crucial parameter to detect sweeps is the window length to compute the statistics that the various methods rely on. The optimal window size depends on the neutral background diversity around the site of interest, which is a consequence not only of the rate of recombination but also the scaled rate of neutral mutations. We choose a constant mutation rate over time, and make the assumption of mutations being introduced during the dormant phase at this constant rate (see equations in introduction). This simplifying assumption is partially supported by empirical evidence (Levin, 1990; Whittle, 2006; Dann et al., 2017), and has so far been made in the wider field of inference models, notably in the ecological sequential Markovian coalescent method (eSMC, Sellinger et al., 2019). While assuming mutation in the dormant population favors the inference of footprints of selection by simply adding additional data, which subsequently increases the likelihood to observe recombination events, it remains unclear if this assumption is justified for all species with a dormant phase and/or if mutations occur at a different rate depending on the age of the dormant population. More research on the rate of mutation and stability of DNA during dormant phases is needed in plant (e.g.Waterworth et al., 2016), fungi and invertebrate species. Nevertheless, even if this mutation rate in seeds is relatively low, our results of a stronger signal of selection under seed banking than in populations without seed banking are still valid. In contrast to the weak seed bank model, it is possible to test for the existence of mutations during the dormant stage under a strong seed bank model as assumed in prokaryotes, because of the much longer dormant phase compared to the coalescence times (Blath et al., 2020).

Finally, as for all sweep models, we show that selective events that are too far back in the past cannot be detected under seed banks. Nonetheless, we show that when there is a seed bank, older sweeps can be detected with increasing accuracy. The presence of a long persistent seed bank could therefore be convenient when studying older adaptation events in plants, fungi and invertebrates that have some form of dormancy. This prediction also agrees with the previous observation that the footprint of older demographic events is stored in the seed bank (predicted in Živković and Tellier, 2012, observed theoretically in Sellinger et al., 2019, and empirically observed in Daphnia in Möst et al., 2015). Our results open avenues for further testing the correlation between past demographic events and selective events for species that present this life-history strategy. However, current methods estimating the age of selective sweeps (Tournebize et al., 2019; Bisschop et al., 2021) would need to use an ad hoc simulator (e.g. such as the one we present here) to generate neutral and selected simulations under seed banking.

### 4.3 Strengths and limitations of the simulation method

The simulation program developed and used in this work, written in C++, is centered on the use of *tskit*. The toolkit allows for the efficient storage of genealogies through time, by removing lineages that have effectively gone extinct in the current population, thus simplifying the genealogy at regular intervals during the program run-time. Despite all our efforts to streamline the process, forward simulations are inherently limited, because each generation has to be produced sequentially. Thus, while being more flexible and intuitively easier to understand than their coalescent counterparts, forward simulations sacrifice computational efficiency in terms of memory and speed. While simulating hundreds or thousands of individuals is possible (also storing their genealogies in a reasonable amount of time), this limitation becomes exaggerated when adding genomic phenomena such as recombination, and even more so when considering ecological characteristics such as seed banking. The latter scales the process of finding the most recent common ancestor by an inverse factor of *b*^2^. As this leads to an increase in run-time of the order of *O*(1*/b*^2^), we kept the population size at 500 (hermaphroditic) diploid individuals. Furthermore, the output format of the simulations are tree sequences, which enables downstream processing and data analysis without the elaborate design of highly specific code. We believe that our code is the first to allow simulations of long stretches of DNA under the seed bank model including recombination and selection. In a previous study, we developed a modified version of the neutral coalescent simulator *scrm* (Staab et al., 2015) which includes a seed bank with recombination (Sellinger et al., 2019). Our current simulator can be used to study the effect and signatures of selection along the genome under dormancy for non-model species with reasonably small population sizes. For a strict application of our model to diploid plants, future work would need to consider the constraint of having only *N* individual diploid parents to choose from. We expect this to likely yield slightly shorter coalescent times than in our pseudo-diploid model (based on the haploid Kaj et al., 2001), while our insights should still be valid.

### 4.4 Towards more complete scenarios of selection

We here explore a scenario in which a single beneficial allele is introduced. The much longer times to fixation in the presence of seed banks suggest that such a scenario may be unlikely. Indeed, it is probable that several alleles under selection, potentially affecting the same biological processes, are maintained simultaneously in populations for longer periods of time. We can therefore surmise that under seed banking, polygenic selective processes and/or competing selective sweeps, often associated with complex phenotypes and adaptation to changing environmental conditions in space and time, should be common.

From the point of view of genomic signatures of selection, the overall effectiveness of selection at a locus coupled with increased effective recombination with seed banking generate narrower selective sweeps, hence less genetic hitch-hiking throughout the genome. While we show that these effects can be advantageous to detect selective sweeps, we speculate that this might not be the case for balancing selection. If seed banks do promote balancing selection (Tellier and Brown, 2009), the expected genomic footprints would be likely narrowly located around the site under selection, and the excess of nucleotide diversity would not be significantly different from the rest of the genome. The presence of seed banking would therefore obscure the signatures of balancing selection. Concomitantly, the Hill-Robertson-Effect and background selection are expected to be weaker under longer seed banks. These predictions could ultimately define the relationship between linkage disequilibrium, the efficacy of selection and observed nucleotide diversity in species with seed banks compared to species without it (Tellier, 2019, Živković and Tellier, 2018).

## Acknowledgements

The authors gratefully acknowledge the computational and data resources provided by the Leibniz Supercomputing Centre (www.lrz.de). KK is supported by a grant from the Deutsche Forschungsgemeinschaft (DFG) through the TUM International Graduate School of Science and Engineering (IGSSE), GSC 81, within the project GENOMIE QADOP. AT receives funding from the Deutsche Forschungsgemeinschaft (DFG) grant TE809/1-4, project 254587930. DAA was a Humboldt Post-Doctoral fellow.

## Conflict of interest disclosure

The authors declare that they have no financial conflict of interest with the content of this article.

## Notes

### Competing Interest Statement

The authors have declared no competing interest.

### Summary of Updates

This version is recommended in PCI Evolutionary Biology

## References

Alachiotis, N. and Pavlidis, P. (2016). Scalable linkage-disequilibrium-based selective sweep detection: a performance guide. GigaScience, 5:7.

Alachiotis, N., Stamatakis, A., and Pavlidis, P. (2012). OmegaPlus: a scalable tool for rapid detection of selective sweeps in whole-genome datasets. Bioinformatics, 28(17):2274–2275.

Barrett, R. D. H., M’Gonigle, L. K., and Otto, S. P. (2006). The Distribution of Beneficial Mutant Effects Under Strong Selection. Genetics, 174(4):2071–2079.

Bisschop, G., Lohse, K., and Setter, D. (2021). Sweeps in time: leveraging the joint distribution of branch lengths. Genetics, 219(2):iyab119.

Blath, J., Buzzoni, E., González Casanova, A., and Wilke-Berenguer, M. (2019). Structural properties of the seed bank and the two island diffusion. Journal of Mathematical Biology, 79(1):369–392.

Blath, J., Buzzoni, E., Koskela, J., and Wilke Berenguer, M. (2020). Statistical tools for seed bank detection. Theoretical Population Biology, 132:1–15.

Blath, J., Casanova, A. G., Kurt, N., and Wilke-Berenguer, M. (2016). A NEW COALESCENT FOR SEED-BANK MODELS. The Annals of Applied Probability, 26(2):857–891.

Blath, J., González Casanova, A., Eldon, B., Kurt, N., and Wilke-Berenguer, M. (2015). Genetic Variability Under the Seedbank Coalescent. Genetics, 200(3):921–934.

Brown, J. H. and Kodric-Brown, A. (1977). Turnover Rates in Insular Biogeography: Effect of Immigration on Extinction. Ecology, 58(2):445–449.

Cohen, D. (1966). Optimizing reproduction in a randomly varying environment. Journal of Theoretical Biology, 12(1):119–129.

Dann, M., Bellot, S., Schepella, S., Schaefer, H., and Tellier, A. (2017). Mutation rates in seeds and seed-banking influence substitution rates across the angiosperm phylogeny. Technical report, bioRxiv. Type: article.

Evans, M. E. K. and Dennehy, J. J. (2005). Germ banking: bet-hedging and variable release from egg and seed dormancy. The Quarterly Review of Biology, 80(4):431–451.

Hairston Jr, N. G. and De Stasio Jr, B. T. (1988). Rate of evolution slowed by a dormant propagule pool. Nature, 336(6196):239–242.

Haller, B. C. and Messer, P. W. (2019). SLiM 3: Forward Genetic Simulations Beyond the Wright-Fisher Model. Molecular Biology and Evolution, 36(3):632–637.

Heinrich, L., Müller, J., Tellier, A., and Živković, D. (2018). Effects of population-and seed bank size fluctuations on neutral evolution and efficacy of natural selection. Theoretical Population Biology, 123:45–69.

Hill, W. G. and Robertson, A. (1968). Linkage disequilibrium in finite populations. TAG. Theoretical and applied genetics. Theoretische und angewandte Genetik, 38(6):226–231.

Hudson, R. R. (1983). Properties of a neutral allele model with intragenic recombination. Theoretical Population Biology, 23(2):183–201.

Kaj, I., Krone, S. M., and Lascoux, M. (2001). Coalescent theory for seed bank models. Journal of Applied Probability, 38:285–300.

Kelleher, J., Thornton, K. R., Ashander, J., and Ralph, P. L. (2018). Efficient pedigree recording for fast population genetics simulation. PLOS Computational Biology, 14(11):e1006581.

Kim, Y. and Stephan, W. (2002). Detecting a local signature of genetic hitchhiking along a recom-bining chromosome. Genetics, 160(2):765–777.

Kimura, M. (1962). On the Probability of Fixation of Mutant Genes in a Population. Genetics, 47(6):713–719.

Koopmann, B., Müller, J., Tellier, A., and Živković, D. (2017). Fisher–Wright model with deter-ministic seed bank and selection. Theoretical Population Biology, 114:29–39.

Lennon, J. T., den Hollander, F., Wilke-Berenguer, M., and Blath, J. (2021). Principles of seed banks and the emergence of complexity from dormancy. Nature Communications, 12(1):4807.

Levin, D. A. (1990). The Seed Bank as a Source of Genetic Novelty in Plants. The American Naturalist, 135(4):563–572.

Manna, F., Pradel, R., Choquet, R., Fréville, H., and Cheptou, P.-O. (2017). Disentangling the role of seed bank and dispersal in plant metapopulation dynamics using patch occupancy surveys. Ecology, 98(10):2662–2672.

Maynard Smith, J. and Haigh, J. (1974). The hitch-hiking effect of a favourable gene. Genetics Research, 23(1):23–35.

Möst, M., Oexle, S., Marková, S., Aidukaite, D., Baumgartner, L., Stich, H.-B., Wessels, M., Martin-Creuzburg, D., and Spaak, P. (2015). Population genetic dynamics of an invasion reconstructed from the sediment egg bank. Molecular Ecology, 24(16):4074–4093.

Nara, K. (2009). Spores of ectomycorrhizal fungi: ecological strategies for germination and dormancy. New Phytologist, 181(2):245–248.

Nei, M. and Li, W. H. (1979). Mathematical model for studying genetic variation in terms of restriction endonucleases. Proceedings of the National Academy of Sciences, 76(10):5269–5273.

Nunney, L. and Ritland, A. E. K. (2002). The Effective Size of Annual Plant Populations: The Interaction of a Seed Bank with Fluctuating Population Size in Maintaining Genetic Variation. The American Naturalist, 160(2):195–204.

Pavlidis, P., Živković, D., Stamatakis, A., and Alachiotis, N. (2013). SweeD: Likelihood-Based Detec-tion of Selective Sweeps in Thousands of Genomes. Molecular Biology and Evolution, 30(9):2224–2234.

Sellinger, T., Abu Awad, D., Möst, M., and Tellier, A. (2019). Inference of past demography, dormancy and self-fertilization rates from whole genome sequence data. preprint, Evolutionary Biology.

Sellinger, T. P. P., Abu-Awad, D., and Tellier, A. (2021). Limits and convergence properties of the sequentially Markovian coalescent. Molecular Ecology Resources, 21(7):2231–2248.

Shoemaker, W. R. and Lennon, J. T. (2018). Evolution with a seed bank: The population genetic consequences of microbial dormancy. Evolutionary Applications, 11(1):60–75.

Shoemaker, W. R., Polezhaeva, E., Givens, K. B., and Lennon, J. T. (2022). Seed banks alter the molecular evolutionary dynamics of Bacillus subtilis. Genetics, 221(2). iyac071.

Staab, P. R., Zhu, S., Metzler, D., and Lunter, G. (2015). scrm: efficiently simulating long sequences using the approximated coalescent with recombination. Bioinformatics, 31(10):1680–1682.

Stephan, W. (2019). Selective Sweeps. Genetics, 211(1):5–13.

Tajima, F. (1983). Evolutionary relationship of DNA sequences in finite populations. Genetics, 105(2):437–460.

Tajima, F. (1989). Statistical method for testing the neutral mutation hypothesis by DNA polymor-phism. Genetics, 123(3):585–595.

Tellier, A. (2019). Persistent seed banking as eco-evolutionary determinant of plant nucleotide diversity: novel population genetics insights. New Phytologist, 221(2):725–730.

Tellier, A. and Brown, J. K. M. (2009). The influence of perenniality and seed banks on polymor-phism in plant-parasite interactions. The American Naturalist, 174(6):769–779.

Tellier, A., Laurent, S. J. Y., Lainer, H., Pavlidis, P., and Stephan, W. (2011). Inference of seed bank parameters in two wild tomato species using ecological and genetic data. Proceedings of the National Academy of Sciences, 108(41):17052–17057.

Templeton, A. R. and Levin, D. A. (1979). Evolutionary Consequences of Seed Pools. The American Naturalist, 114(2):232–249.

Tournebize, R., Poncet, V., Jakobsson, M., Vigouroux, Y., and Manel, S. (2019). McSwan: A joint site frequency spectrum method to detect and date selective sweeps across multiple population genomes. Molecular Ecology Resources, 19(1):283–295.

Verin, M. and Tellier, A. (2018). Host-parasite coevolution can promote the evolution of seed banking as a bet-hedging strategy. Evolution, 72(7):1362–1372.

Vitalis, R., Glémin, S., and Olivieri, I. (2004). When genes go to sleep: the population genetic conse-quences of seed dormancy and monocarpic perenniality. The American Naturalist, 163(2):295–311.

Waterworth, W. M., Footitt, S., Bray, C. M., Finch-Savage, W. E., and West, C. E. (2016). DNA damage checkpoint kinase ATM regulates germination and maintains genome stability in seeds. Proceedings of the National Academy of Sciences of the United States of America, 113(34):9647–9652.

Whittle, C.-A. (2006). The influence of environmental factors, the pollen : ovule ratio and seed bank persistence on molecular evolutionary rates in plants. Journal of Evolutionary Biology, 19(1):302–308.

Willis, C. G., Baskin, C. C., Baskin, J. M., Auld, J. R., Venable, D. L., Cavender-Bares, J., Donohue, K., Rubio de Casas, R., and NESCent Germination Working Group (2014). The evolution of seed dormancy: environmental cues, evolutionary hubs, and diversification of the seed plants. The New Phytologist, 203(1):300–309.

Živković, D. and Tellier, A. (2012). Germ banks affect the inference of past demographic events. Molecular Ecology, 21(22):5434–5446.

Živković, D. and Tellier, A. (2018). All But Sleeping? Consequences of Soil Seed Banks on Neutral and Selective Diversity in Plant Species. In Morris, R. J., editor, Mathematical Modelling in Plant Biology, pages 195–212. Springer International Publishing, Cham.

