## Supplementary material for "Weak seed banks influence the signature and detectability of selective sweeps": SI Appendix

Table S1: Proposed number of generations to simulate before adding selective mutation (for 2N=1000). Also, the number of generations simulated to estimate TMRCA.

| germination rate (b) | recombination rate (r) | calibration generations |
|---|---|---|
| 1 | 0 | 40000 |
| 0.5 | 0 | 80000 |
| 0.35 | 0 | 160000 |
| 0.25 | 0 | 320000 |
| 1 | $10^{-8}$ | 48000 |
| 0.5 | $10^{-8}$ | 96000 |
| 0.35 | $10^{-8}$ | 192000 |
| 0.25 | $10^{-8}$ | 384000 |
| 1 | $10^{-7}$ | 56000 |
| 0.5 | $10^{-7}$ | 112000 |
| 0.35 | $10^{-7}$ | 224000 |
| 0.25 | $10^{-7}$ | 448000 |
| 1 | $10^{-6}$ | 64000 |
| 0.5 | $10^{-6}$ | 128000 |
| 0.35 | $10^{-6}$ | 256000 |
| 0.25 | $10^{-6}$ | 512000 |

### A Absolute TMRCAs for different germination and recombination rates

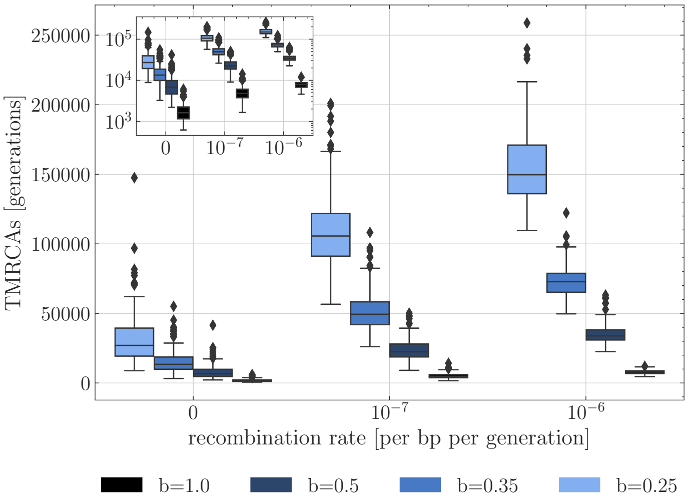


Figure S1: Absolute time to the most recent common ancestor (TMRCA) as a function of the germination rate $b$. For each germination rate, three recombination rates per site are presented ($r = 0$, $r = 10^{-7}$ and $r = 10^{-6}$. Boxes describe the 25th (Q1) to 75th percentile (Q3), with the lower whisker representing Q1-1.5×(Q3-Q1) outlier threshold and the upper whisker is calculated analogously. The mean is plotted between Q3 and Q1. Each boxplot represents the distribution of 200 TMRCA values over 200 sequences of 0.1 Mb. Per sequence the oldest TMRCA is retained.

### B Fixation time phase contribution

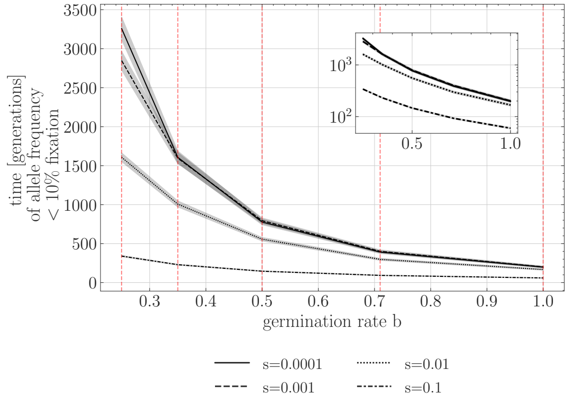


(a)

Figure S2: Time to fixation for different selection coefficients. Y-axis is the unnormalized time in generations, and X-axis is the germination rate $b$. a) Time allele spends below 10% frequency in population and b) time allele spends above 10 % and below 90% frequency c) above 90% in population. A dormancy effective population size coefficient $N_e^b s$ can be calculated with each intersection of vertical dashed lines with fixation times by scaling with $b^2$, e.g. for $N_e^{b=1.0} s = 1$: $N_e^{b=0.71} s = 2.0$, $N_e^{b=0.5} s = 4$, $N_e^{b=0.35} s = 8.2$, $N_e^{b=0.25} s = 16$. In total 1000 replicates were used for each parameter configurations

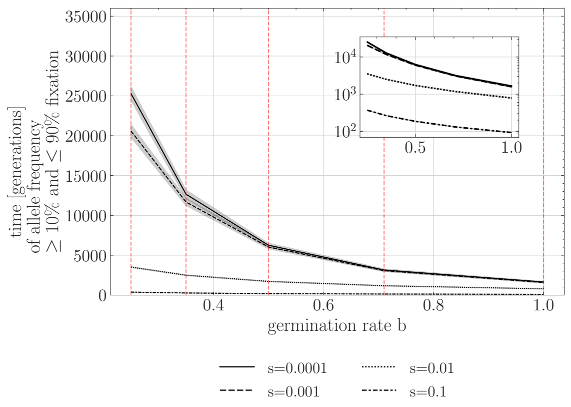


(b)

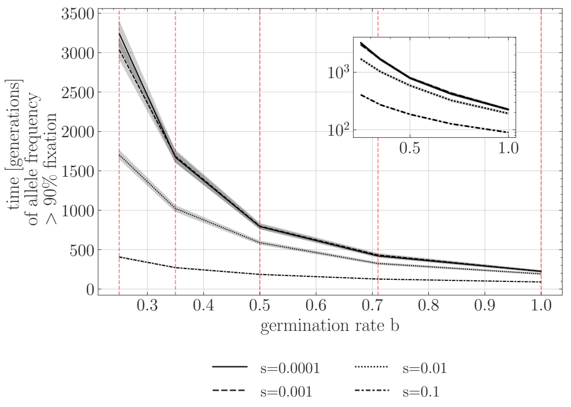


(c)

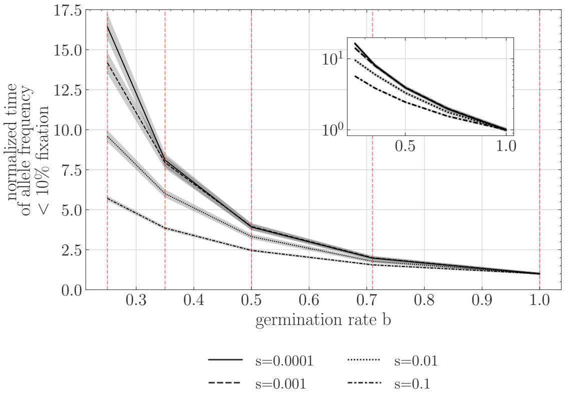


(a)

Figure S3: Time to fixation for different selection coefficients. Y-axis is the time normalized by the $b = 1$ estimate for each respective selection coefficient, and X-axis is the germination rate $b$. a) Time allele spends below 10% frequency , b) time allele spends above or equal to 10 % and below or equal to 90% frequency and c) above 90% frequency in population. A dormancy effective population size coefficient $N_e^b s$ can be calculated with each intersection of vertical dashed lines with fixation times by scaling with $b^2$, e.g. for $N_e^{b=1.0}s = 1$: $N_e^{b=0.71}s = 2.0$, $N_e^{b=0.5}s = 4$, $N_e^{b=0.35}s = 8.2$, $N_e^{b=0.25}s = 16$. In total 1000 replicates were used for each parameter configurations

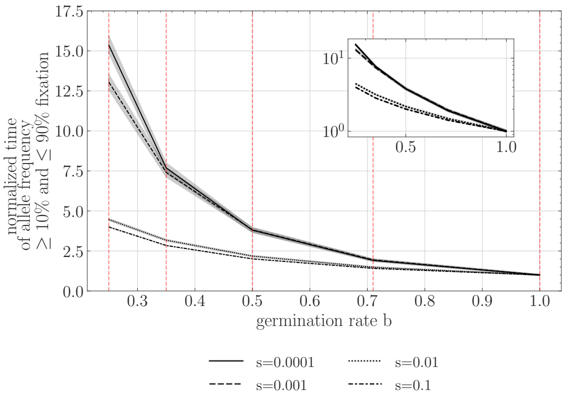


(b)

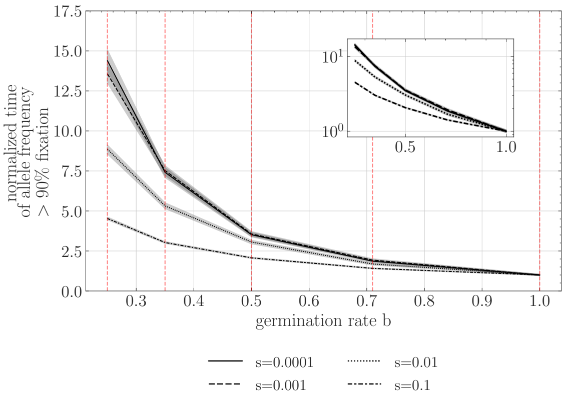


(c)

Figure S4: Time contribution to fixation for different selection coefficients in percent of the phases (a) below 10 % allele frequency and (b) above 10 % and below 90% allele frequency and (c) above 90% allele frequency. In total 1000 replicates were used for each parameter configurations.

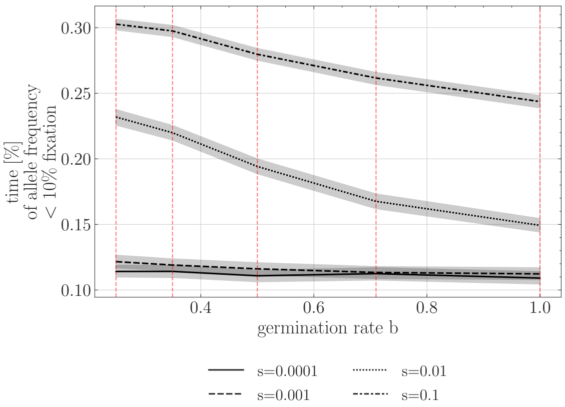


(a)

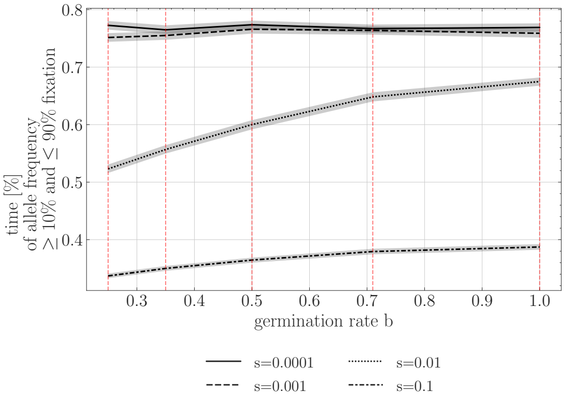


(b)

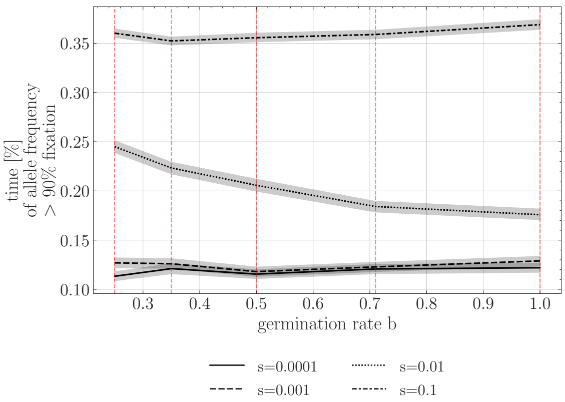


(c)

### 4 C Sweep recovery signatures after fixation

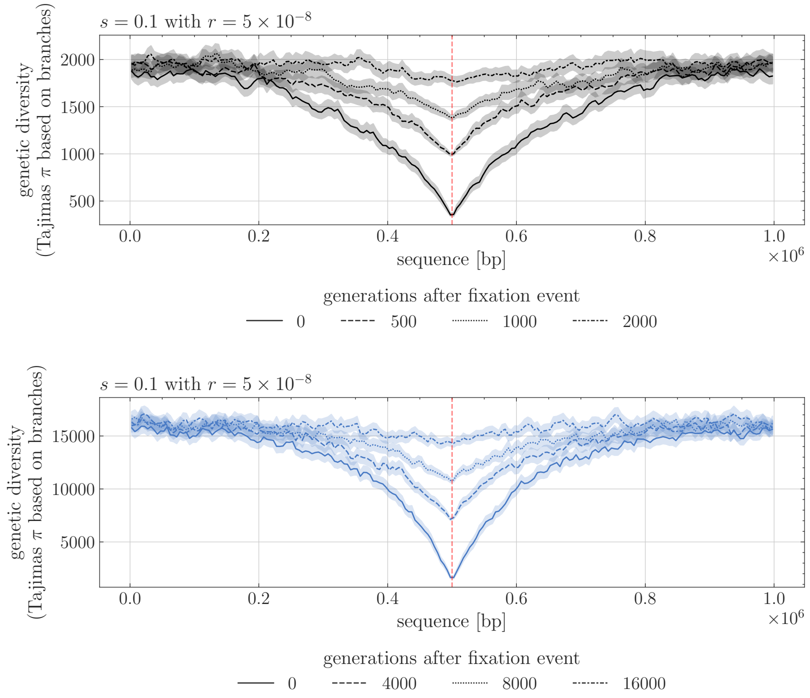


Figure S5: Nucleotide diversity (Tajima's $\pi$, Y-axis) over sequence length (X-axis) for windows of size 5000 mapping length and averaged over 400 repetitions. Comparison between germination rate a) $b = 1$ and b) $b = 0.35$ for different sweep recovery times. A selection coefficient of $N_e^{b=1} s = 100.0$ and recombination rate $r = 5 \times 10^{-8}$ per generations per bp was set for all simulations.

### D Effect of different dominance coefficients

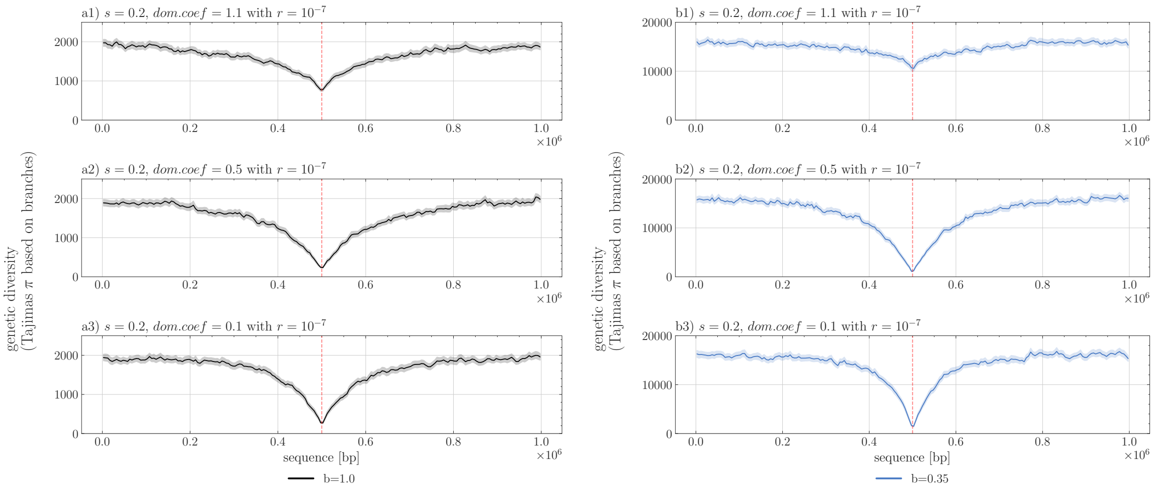


Figure S6: Nucleotide diversity (Tajima's $\pi$, Y-axis) over sequence length (X-axis) for windows of size 5000 and averaged over 200 repetitions. Comparison between germination rate a) $b = 1$ and b) $b = 0.35$ for different dominance coefficients a1, b1) $h = 1.1$, a2,b2) $h = 0.5$, a3, b3) $h = 0.1$, respectively. A selection coefficient of $N_e^{b=1}s = 200$ and recombination rate $r = 5 \times 10^{-7}$ per bp per generation was set for all simulations.

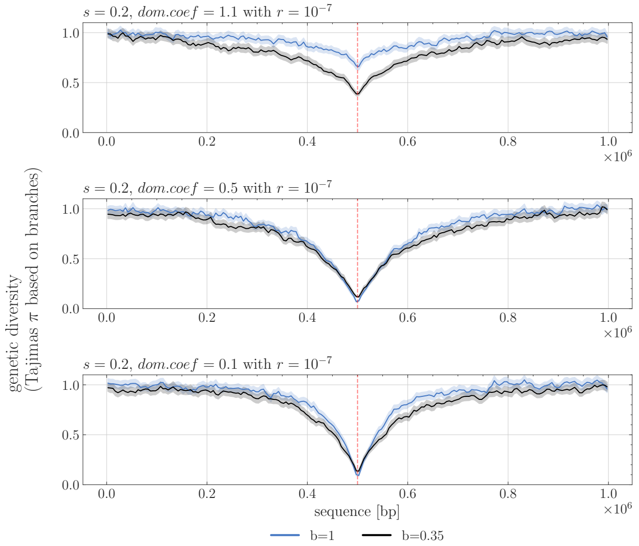


Figure S7: Normalized nucleotide diversity (Tajima's $\pi$, Y-axis) over sequence length (X-axis) for windows of size 5000 and averaged over 200 repetitions. Comparison between germination rate $b = 1$ and $b = 0.35$ for different dominance coefficients $h = 1.1$, $h = 0.5$, $h = 0.1$, respectively. A selection coefficient of $N_e^{b=1}s = 200$ and recombination rate $r = 5 \times 10^{-7}$ per bp per generation was set for all simulations.

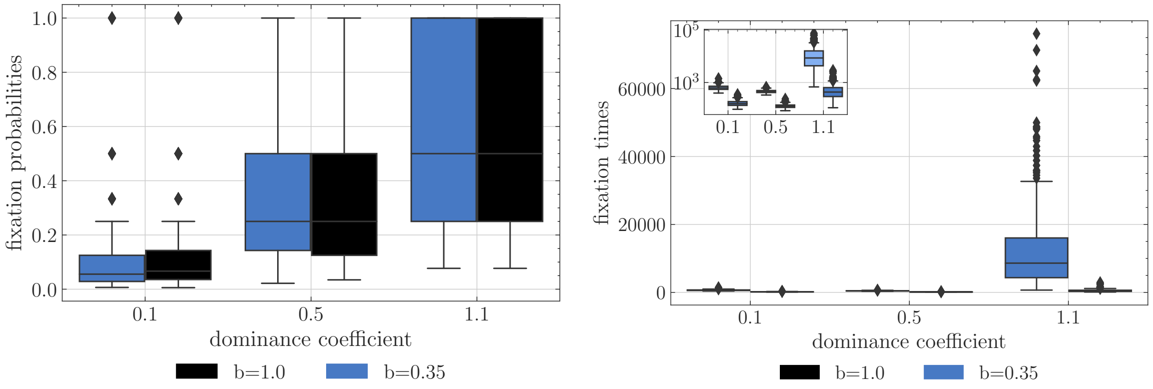


(a) (b)

Figure S8: Fixation probability (a) and time (b) for different dominance coefficients and two different germination rates, namely $b = 1$ and $b = 0.35$ for 400 replicates. Selection coefficient was set to 0.2, corresponding to $N_e^{b=1}s = 200$ and $N_e^{b=0.35}s = 1632.7$. In total 1000 replicates were used for each parameter configurations and simulations were conditioned on fixation.

### E Scaling population size by $\frac{1}{b^2}$

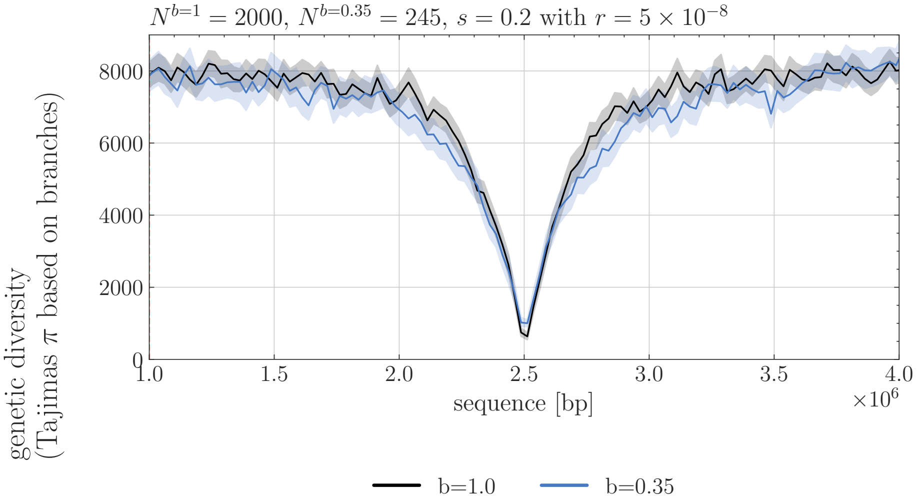


Figure S9: Tajima's $\pi$, Y-axis) over sequence length of 5 mb (X-axis) for windows of size 25000 and averaged over 150 repetitions. Comparison between germination rate $b = 1$ and $b = 0.35$ under a selection coefficient of $s = 0.2$, corresponding to $N_e^{b=1}s = 400$ without a seed bank (black) and to $N_e^{b=0.35}s = 400$ (blue), assuming population sizes of 2000 and 245 diploid individuals, for no seed bank and seed bank, respectively. A recombination rate of $r = 5 \times 10^{-8}$ per bp per generation was set for all simulations.

### F Narrow sweep signature of a large sequence lengths

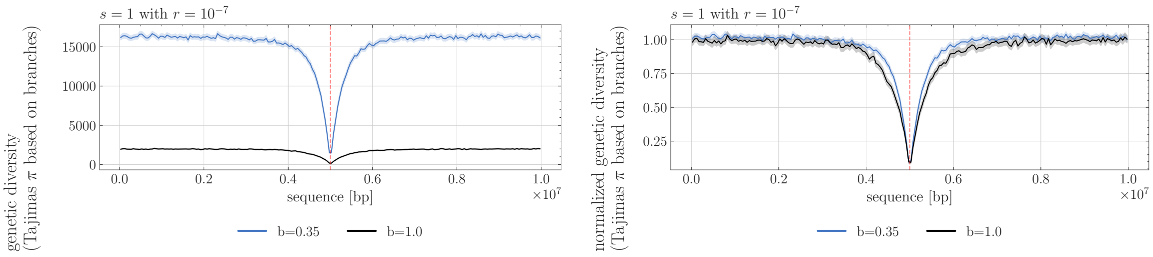


(a) (b)

Figure S10: Nucleotide diversity (a) Tajimas $\pi$ and (b) normalized diversity Y-axis) over sequence length of 10 mb (X-axis) for windows of size 50000 and averaged over 400 repetitions, the shaded area represents a 95% confidence interval. Comparison between germination rate $b = 1$ (black) and $b = 0.35$ (blue) for selection coefficients $s = 1$ corresponding to $N_e^{b=1}s = 1000$ and $N_e^{b=0.35}s = 8163.3$ with a dominance coefficient of $h = 0.5$ and recombination rate of $r = 10^{-7}$ per bp and generation.
